# *In Silico*-Driven Engineering of *Halomonas elongata* L-Asparaginase: Towards Enhanced Proteolytic Resistance in Lymphoblastic Leukemia

**DOI:** 10.1101/2024.06.07.597648

**Authors:** Maryam Samadaei Ghadikolaei, Sedigheh Asad, Vahideh Hassan-Zadeh

## Abstract

The shortened L-asparaginase’s half-life in leukemia patients due to elevated serum proteases, poses a challenge. This study aimed to enhance the stability of *Halomonas elongata* L-asparaginase against trypsin. Employing the trRosetta server, we modeled the enzyme’s 3D structure with a quality score of 96.5, revealing predominant secondary structure of random coils (42%), alpha helices (33%), and extended strands (20%) organized in two domains. Molecular docking unveiled a triad alignment among residues Thr16, Ser65, and Asp97 with L-asparagine. Site selection for mutation considered secondary structure prediction, dimerization analysis, trypsin cleavage site determination and epitope mapping. A library of enzyme variants was constructed through site saturation mutagenesis which led to the identification of the Arg206 to Thr, resulting in a 1.7-fold increased enzyme-specific activity (2400 U/mg) and heightened trypsin resistance. The mutant displayed a half-life of 3.47 hin human serum, approximately 50% longer than the wild type. *In silico* analyses confirmed structural stability, reduced flexibility, and enhanced substrate binding, contributing to increased proteolysis resistance and enzymatic activity. The Arg206Thr mutant exhibited anti-proliferative activity (IC_50_ of 1.45 U/ml) on leukemia cell line K562, suggesting potential therapeutic implications.

## 1 Introduction

Acute lymphoblastic leukemia (ALL) constitutes a malignancy primarily affecting bone marrow, representing a significant cause of mortality in pediatric populations [1]. L-asparaginase (L-ASNase) (EC 3.5.1.1) emerges as a pivotal therapeutic agent in ALL treatment [2]. This particular enzyme holds significant importance as it is the first bacterial enzyme utilized in this treatment. Its primary function involves catalyzing the hydrolysis of the L-asparagine (L-Asn) into L-aspartate (L-Asp) and ammonia [3]. The efficacy of L-ASNase lies in its disruption of protein synthesis within malignant cells, exploiting their dependency on extracellular L-Asn sources, while normal cells possess endogenous L-Asn synthesis pathways [4]. Bacterial L-ASNases, such as those from *Escherichia coli* and *Erwinia chrysanthemi*, are generally utilized in clinical treatment [5, 6]. However, due to their bacterial origin, can elicit immune responses and trigger the production of antibodies, which can further diminish its efficacy [7, 8]. Consequently, repeated administration is required to maintain therapeutic levels, resulting in the development of anti-drug antibodies (ADAs) [9, 10]. Certain ALL patients, particularly those with malignant subtypes, exhibit diminished response to L-ASNase therapy due to heightened lysosomal cysteine protease expression, notably asparaginyl endopeptidase (AEP) and cathepsin B [11, 12]. The action of these proteases leads to the cleavage of L-ASNase, rendering it inactive, and the subsequent antigenic epitopes exposure that trigger allergic reactions. Additionally, trypsin-mediated proteolytic degradation further compromises L-ASNase stability and half-life in the bloodstream, requiring frequent dosing [13].

One strategy to enhance the stability and decrease the immune response of L-ASNase involves the covalent binding of the enzyme to polyethylene glycol (PEG); however, this method may also lead to adverse reactions [14, 15]. Current investigations are focused on alternative approaches to prolong the half-life of L-ASNase within the circulatory system, including the utilization of non-immunogenic and biodegradable unstructured polymers like hydrophilic protein-polymer (XTEN), Pro-Ala and Ser (PASylation), encapsulation within red blood cells (RBC), and the development of novel enzyme formulations [16, 18]. In addition, novel microbial sources of L-ASNase are currently under investigation, aiming to identify those with distinct characteristics. Another strategy employed to enhance the performance of L-ASNase involves protein engineering, which focuses on improving its thermodynamic stability, substrate specificity, catalytic activity and solubility. Protein engineering, utilizing rational design, directed evolution, or a combination of both methods, plays a pivotal role in augmenting the properties of proteins [19, 21]. Notably, protein engineering has demonstrated success in enhancing the stability, enzyme activity, and immunogenicity of L-ASNase variants [22, 25].

Before, a novel L-ASNase derived from the halophilic bacterium *Halomonas elongata* exhibited remarkable specificity towards L-Asn and cytotoxicity against leukemia cell lines [26]. Leveraging computational tools for structural modeling and mutational analysis, we aim to identify stability-conferring mutants for enhancing the protease resistance of this L-ASNase in the blood serum. The experimental methodology involves site-saturation mutagenesis, recombinant expression in *E. coli* BL21, and subsequent evaluation of proteolytic stability, enzymatic activity, and anticancer efficacy of both wild-type and mutant L-ASNases.

## 2 Experimental Procedures

### 2.1. Chemicals, bacterial strains, and cell lines

Primers and DNA ladder were ordered from SinaClon (Iran). High-Fidelity DNA polymerase and components of PCR reaction were from kawsarbiotech (Iran). Molecular biology enzymes were purchased from Thermo Fischer (USA). Isopropyl-β-D thiogalactopyranoside (IPTG) and trypsin were from Sigma-Aldrich (USA). *E. coli* XJb (DE3) strain was from Zymo Research (USA). *H. elongata* IBRCM10216 (DSM 2581T), human umbilical vein endothelial cells (HUVEC), and Human Chronic Myelogenous Leukemia (K562) were purchased from Iranian Biological Resource Center (IBRC). The chemicals utilized were of high purity grade suitable for analytical purposes.

### 2.2. Three-Dimensional Protein Modeling and Structural Studies

The study initiated by retrieving the nucleotide sequences of the *asn* gene of *H. elongata* from GenBank accession number KY508642.1. Subsequently, a BLASTn search was conducted to identify suitable templates for comparative or homology modeling, utilizing sequences from the NCBI database. Following this, multiple sequence alignment analysis was done using PRALINE (http://www.ibi.vu.nl/programs/pralinewww/), comparing the *H. elongata* L-ASNase protein sequence with others available in databases. The PSIPRED server (http://bioinf.cs.ucl.ac.uk/psipred/) was employed for the prediction of the enzyme’s secondary structure, offering insights into the positioning of α-helices, β-strands, turns, and random coils within its structure. Structural modeling of *H. elongata* L-ASNase was achieved through the trRosetta server (https://yanglab.nankai.edu.cn/trRosetta/), which integrated deep learning and homologous templates. Among the models generated, the one with the highest TM score, indicative of accurately predicted topology, was selected for further analysis. To assess the stereochemical quality of the modeled L-ASNase, PROCHECK (http://www.ebi.ac.uk/thornton-srv/software/PROCHECK/) and the VADAR (Volume Area Dihedral Angle Reporter) servers (http://redpoll.pharmacy.ualberta.ca/vadar.) were utilized. Additionally, alignments of the structure and sequence were created, and the global root-mean-square deviation (RMSD) measurements between the template and target were computed using a superpose server. Finally, an analysis of different types of atoms and non-bonded interactions in the refined model was carried out using the ERRAT server.

In the process of identifying domains, dimerization, and epitope mapping, the Protein Peeling 2 (PP2) online server was employed to predict protein units within the modeled enzyme. PP2 operates based on a contact probability matrix. The quaternary protein structure was predicted via the COTH (CO-THreader) online server (https://zhanggroup.org/COTH/), employing template-based modeling techniques. Subsequently, the residues involved in the formation of the quaternary structure were identified using PDBePISA (https://www.ebi.ac.uk/pdbe/api/doc/pisa.html/). For the docking of the ligand L-Asn to the L-ASNase enzyme, the active site residues of the modeled L-ASNase were anticipated using multiple sequence alignments and various online servers such as InterProScan (http://www.ebi.ac.uk/InterProScan/), PSI-BLAST (https://www.ncbi.nlm.nih.gov/BLAST/), and Galaxy servers (http://galaxy.seoklab.org/site). The ligand L-Asn (with a molecular mass of 132.12 g/mol) was obtained in SDF format from the PubChem database. It was then docked onto the catalytic site of *H. elongata* L-ASNase using the SwissDock web server (https://www.swissdock.ch/). Visualization of the models and the docked complexes was performed using PyMoL.

### 2.3. Development of Mutations and Smart Libraries Guided by Sequence Input Data

Predictions for cleavage sites within the protein sequence were made using the PeptideCutter (https://web.expasy.org/peptide_cutter/), while the Porter 4.0-PaleAle 4.0 was utilized to forecast the accessible surface area of these cleavage sites. Further analysis involved the use of the LBtope web server (https://webs.iiitd.edu.in/raghava/lbtope/) to predict B-cell epitopes. Mutagenesis hotspots within the modeled *H. elongata* L-ASNase were discerned employing the HotSpot Wizard2 (http://loschmidt.chemi.muni.cz/hotspotwizard). Subsequent to hotspot identification, the DynaMut server (http://biosig.unimelb.edu.au/dynamut/) was harnessed to forecast the impact of substitutions on the thermodynamic stability of proteins, additionally, the Residue Interaction Network Generator (RING) (https://ring.biocomputingup.it/) was employed to explore diverse intra-atomic bindings present in L-ASNase, encompassing ionic bonds, hydrogen bonds, disulfide bonds, π-cation interactions, and π-π stacking interactions.

### 2.4. Saturation Mutagenesis and Library Screening

The coding sequence of the *asp* gene, previously cloned in the *XhoI*-*NdeI* sites of the pET-21a vector, served as the template for performing mutagenesis. Site-saturated mutagenesis was conducted using back-to-back primers, adhering to the principles of the Q5 site directed mutagenesis method with minor adjustments [27]. The forward primer, incorporating the degenerate codon NNK at the targeted site, and the 5’-phosphorylated reverse primer were as follows: 5’-GCTTGGAACGCCAGNNKAGCGCACCTCGCTT-3’ and 5’p-CACGACCCGGAAACAGGACAGCGTCGTCGTCAACG-3’, respectively.

The purified PCR products were self-ligated, and transformed into the *E. coli* XJb (DE3). Transformation mixture was spread on Luria-Bertani (LB) agar plates (100 μg/mL ampicillin) and grown at 37 °C overnight. Later, the transformed colonies were inoculated into 96-well plates (200 μL of LB containing 100 μg/mL ampicillin) and incubated overnight at 37 °C and 150 rpm. Negative and positive controls were employed in this study, namely *E. coli* XJb - pET21a and *E. coli* XJb -pET21a+asn cells, respectively. The creation of the replica plate for storage involved the addition of 100 μL of each culture to 100 μL of 30% glycerol. To initiate enzyme expression, a fresh 96-well plate was prepared by inoculating 5 μL of precultures into 200 μL of LB medium supplemented with 100 μg/mL ampicillin. L-arabinose (3 mM) was added to the culture media to induce the expression of the bacteriophage λ endolysin. Once the OD600 of the culture get to 0.5–0.6, L-ASNase expression was triggered by adding IPTG (0.5 mM), followed by a 16h incubation at 20°C (150 rpm). Subsequently, the cells were collected and preserved at −20°C for a duration of 24h. The combination of endolysin with a single freeze-thaw cycle proved efficient in causing cell lysis within a 10min duration at 37°C. In the following, the lysates were re-suspended in ice-cold lysis buffer (phosphate buffer, 50 mM; NaCl, 300 mM at pH 7.50) and subjected to another round of centrifugation at 9000 g for 30min. The activities of crude enzymes, as well as their residual activities after treatment at 37°C (5min) were then assessed. Mutants showing heightened or comparable specific activities to the wild-type were selected for further analysis. The mutant plasmids were sequenced using the T7 terminator primer (Macrogen, Seoul, Korea).

### 2.5. Characterization of The Selected Mutant

*E. coli* BL21 (DE3) cells were transformed with the plasmids coding selected mutant. IPTG 0.5 mM was used for protein expression induction in inoculated cultures. The bacteria were further grown for a duration of 16h at 20°C, followed by collection through centrifugation at (9000g, 10min at 4°C). The cells were suspended again in cold lysis buffer and disrupted through sonication involving a series of recurring cycles. Each cycle consisted of 10s of pulsing followed by 5s of resting, and this process was repeated for a duration of 10min (SYCLON Ultra Sonic Cell Sonicator SKL950-IIDN). The lysate underwent centrifugation at 9000 g for 30min at 4°C and the resulting supernatant was applied onto a Ni-NTA agarose column that had been pre-equilibrated with lysis buffer. The bound enzyme was eluted by the potassium phosphate buffer supplemented with imidazole (0.2 M at pH 7.5). Purified fractions were subjected to buffer exchange through an Amicon Ultra-15 centrifugal filter unit (Merck Millipore) to eliminate imidazole. All the protein purification procedures were conducted at a temperature of 4°C. To ensure the purity of the enzyme, SDS-PAGE (12% w/v acrylamide) was employed. The gels were then stained with Coomassie Brilliant Blue R-250 (Bio-Rad, USA). The protein concentration was determined using the Bradford method, with bovine serum albumin serving as the standard [28].

To calculate the activity of the L-ASNase, purified enzymes were incubated with L-Asn (12 mM) and phosphate buffer 50 mM (pH 8) at 37 °C for 1min. The amount of ammonia released was calculated using Nessler’s reagent and measured spectrophotometrically at 410 nm using a Biotek Micro Plate Reader (USA) [29]. Various concentrations of ammonium sulfate ((NH_4_)_2_SO_4_) were employed as the standard solutions. To quantify enzyme activity, a single unit (U) was defined as the amount of enzyme that induces the generation of 1 μmole of ammonia within 1min under the given assay conditions. The specific activity was denoted as U/mg protein (μmol.min^-1^. mg^-1^). Various concentrations of L-ASN was used to determine kinetic parameters (1 to 12 mM) with 1.5 μg of enzyme [30]. The Michaelis-Menten equation was employed to fit the data using GraphPad Prism software version 9, resulting in the determination of apparent kinetic values. All experimental procedures were conducted in triplicate, and the mean values accompanied by their corresponding standard deviations (SD) were reported.

### 2.6. Effect of Temperature, pH, and NaCl Concentration on Enzyme Activity

The optimal temperature for enzyme activity was identified by performing the reaction at a range of temperatures, from 25 °C to 65 °C. To find the optimum pH, a buffer containing Tris, phosphate, and glycine (50 mM) ranging from 4 to 9 was prepared [31]. The effect of NaCl on enzyme activity was investigated by subjecting different concentrations of NaCl, varying from 0 to 4 M, to thorough examination. It is noteworthy that the enzyme concentration in all reactions remained constant at 30 μg/mL.

### 2.7. Enzymes Stability Assessment Towards Trypsin and Blood Serum

Proteolysis was carried out in vitro by using trypsin in a solution composed of 10 mM phosphate buffer and 1 mM CaCl_2_ (pH 8), to purified wild-type and mutated L-ASNase at varying ratios of 0.5, 1, 5, and 20%. The concentration of both mutant and wild-type enzymes was maintained at 20 μg/mL. The mixture was placed at 37 °C for varying durations ranging from 0 to 120min. Every 30min, 0.1 mL of the sample was promptly combined with a 24 mM L-Asn solution and subjected to incubation at 37 °C for one min to determine the residual activity.

In vitro experiments were carried out to analyze how serum components and inhibitors could affect the activity and stability of enzymes. This involved mixing the enzyme with both blood serum and phosphate buffer (supplemented with 140 mM NaCl, pH 7.50) concurrently [32]. To assess the half-life of both mutant and wild-type enzymes in blood serum, the enzyme solution was introduced to blood serum at a 1:5 ratios. This resulted in a final concentration of 30 μg/mL of the enzymes in the serum. Subsequently, the tubes were subjected to incubation at 37°C for 28 h, with the residual activity being calculated at regular time intervals.

### 2.8. Enzyme Cytotoxicity Assessment

The K562 cell line was utilized to assess the antitumor activity and inhibitory concentration (IC_50_) of *H. elongata* L-ASNase and its mutated form. The cells were incubated in RPMI 1640 medium containing fetal bovine serum and penicillin before being transferred to a 96-well plate. Following a 24 period, they were exposed to varying levels of wild-type and mutant L-ASNase (0, 0.25, 0.5, 1, 1.75, and 2.5 U/mL) and subsequently incubated for an additional 24h [33]. The MTT (3-(4, 5-dimethylthiazol-2-yl)-2, 5 diphenyltetrazolium bromide) test was employed to measure viability, comparing treated cells to controls treated with phosphate buffered saline [34].

To evaluate potential cytotoxic effects on non-leukemic cells, human umbilical vein endothelial cells (HUVEC) were exposed to different concentrations of wild-type and mutant L-ASNase. Additionally, an in vitro hemolysis test was conducted using heparinized human blood cells incubated at 37°C with different enzyme concentrations (0 to 6 U/mL) for 16 h. The measurement of the supernatant’s absorbance was conducted at a wavelength of 540 nm, comparing against negative controls (blood cells without enzyme) and positive controls (blood cells treated with SDS) [33]. All experiments were performed in triplicates

## 3 Results

### 3.1. Three-Dimensional Protein Modeling and Structural Studies

The monomeric subunit of *H. elongata* L-ASNase consists of 355 amino acids. BLASTn results demonstrated the highest identity with L-ASNase isolated from *Yersinia pestis* (PDB: 3NTX) and the type I L-ASNase from *E. coli* (PDB: 3C17), 37% (53% similarity) and 37.1% (52.6% similarity). Multiple sequences alignment was conducted with six L-ASNase with the highest scores for alignments, including *E. coli* L-ASNase I (PDB: 2P2N), *E. coli* K-12 L-ASNase I (T162A) (PDB: 6NXC), *Y. pestis* L-ASNase I (PDB: 3NTX), *guinea pig* L-ASNase I (PDB: 5DNC), and *E. coli* L-ASNase II (PDB: 3ECA). Among these alignments, only 157 amino acids (22.5%) exhibited a conservation score above 7. However, the presence of conserved motifs across various L-ASNases was confirmed through their degree of correlation (Fig. 1).

**Fig. 1.**
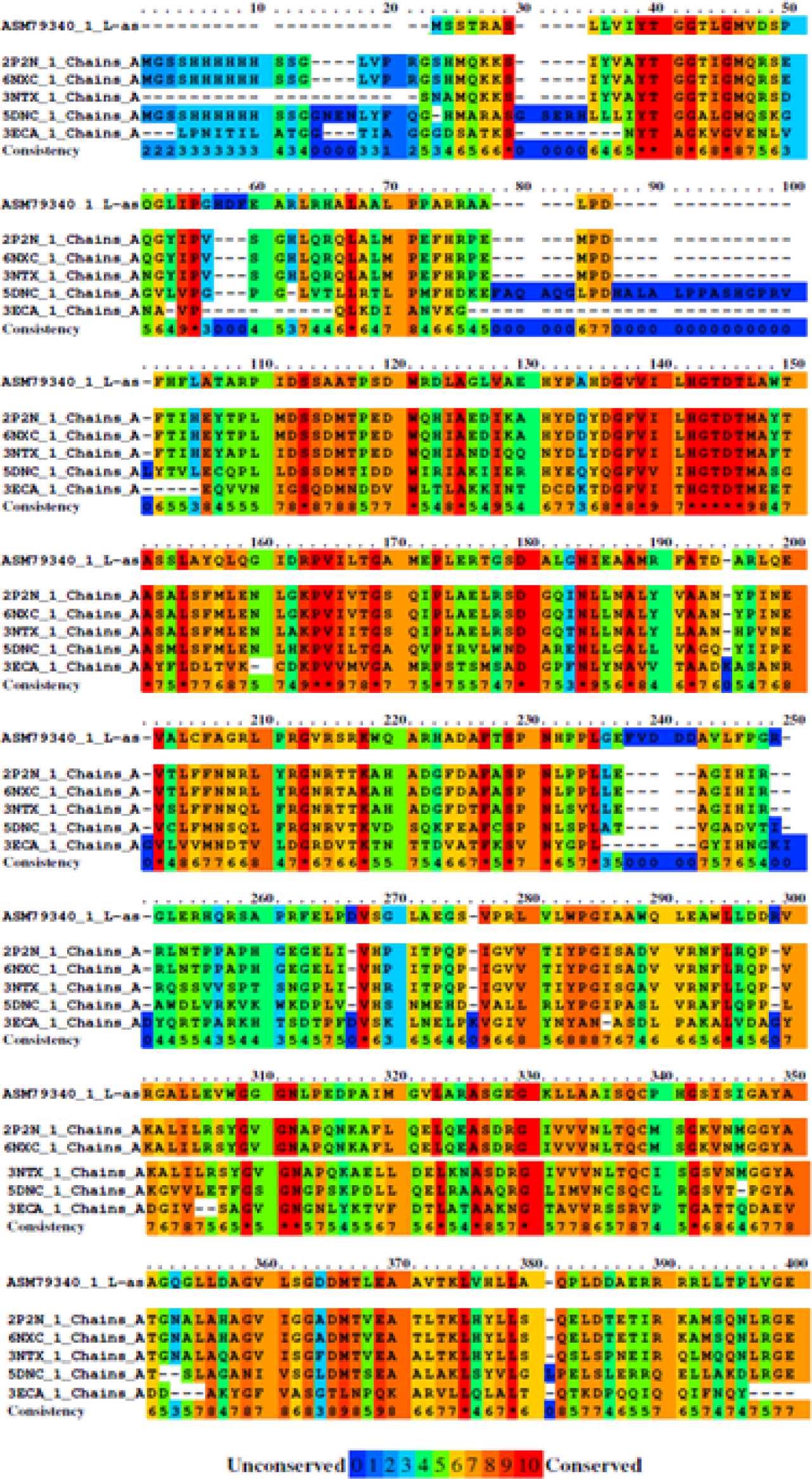
Multiple sequence alignment of *H. elongata* L-ASNase with other L-ASNase sequences exhibiting high alignment scores. Columns highlighted in red indicate residues that are conserved across all sequences. ASM79340.1 : *H. elongata* L-ASNase*, E. coli* K-12 L-ASNase I (T162A) (PDB: 6NXC), *Y. pestis* L-L-ASNase I (PDB: 3NTX), *guinea pig* L-ASNase I (PDB: 5DNC), *E. coli* L-ASNase I (PDB: 2P2N), and *E. coli* L-ASNase II (PDB: 3ECA).

The tertiary structure model of *H. elongata* L-ASNase was predicted, leveraging the 3D structure of *Y. pestis* (PDB: 3NTX) as a reference. Employing template-based modeling (TBM), the trRosetta generated five models; the model exhibiting the highest TM score (0.98) was picked for further analysis due to its demonstrated accuracy and confidence (Fig. 2A, B). The root-mean-square deviation (RMSD) between the modeled enzyme and *Y. pestis* (PDB: 3NTX) was computed, revealing a value of 3.14 when aligning 315 α carbon atoms (Fig. S1A). Analysis of the Φ and Ψ distributions via the Ramachandran plot indicated that 92% of residues reside within the most favored regions, with 5.5% in allowed regions, and negligible proportions in generously allowed or disallowed regions (Fig. S1 B). Thus, all amino acid residues fall within permissible regions, affirming the modeled protein structure’s satisfactory geometric fitness and stereochemistry. ERRAT analysis employed to evaluate the overall quality of the 3D model of the L-ASNase protein, yielded a value of 97.05%, surpassing the rejection threshold of 95% (Fig. S2). The quaternary structure of the enzyme was forecasted through query-template alignment sequences, revealing a dimeric arrangement. Around 50 amino acids play a pivotal role in the dimerization process, establishing connections between two monomers via hydrogen bonds and salt bridges (Fig. 2C). L-ASNase exhibits a secondary structure that is predominantly characterized by random coils (42%), alpha helices (33%), and extended strands (20%). These structural elements are organized into two distinct domains: a larger N-terminal domain and a smaller C-terminal domain which are connected by approximately 15 residues (Fig. 2 and Fig.3).

**Fig.2.**
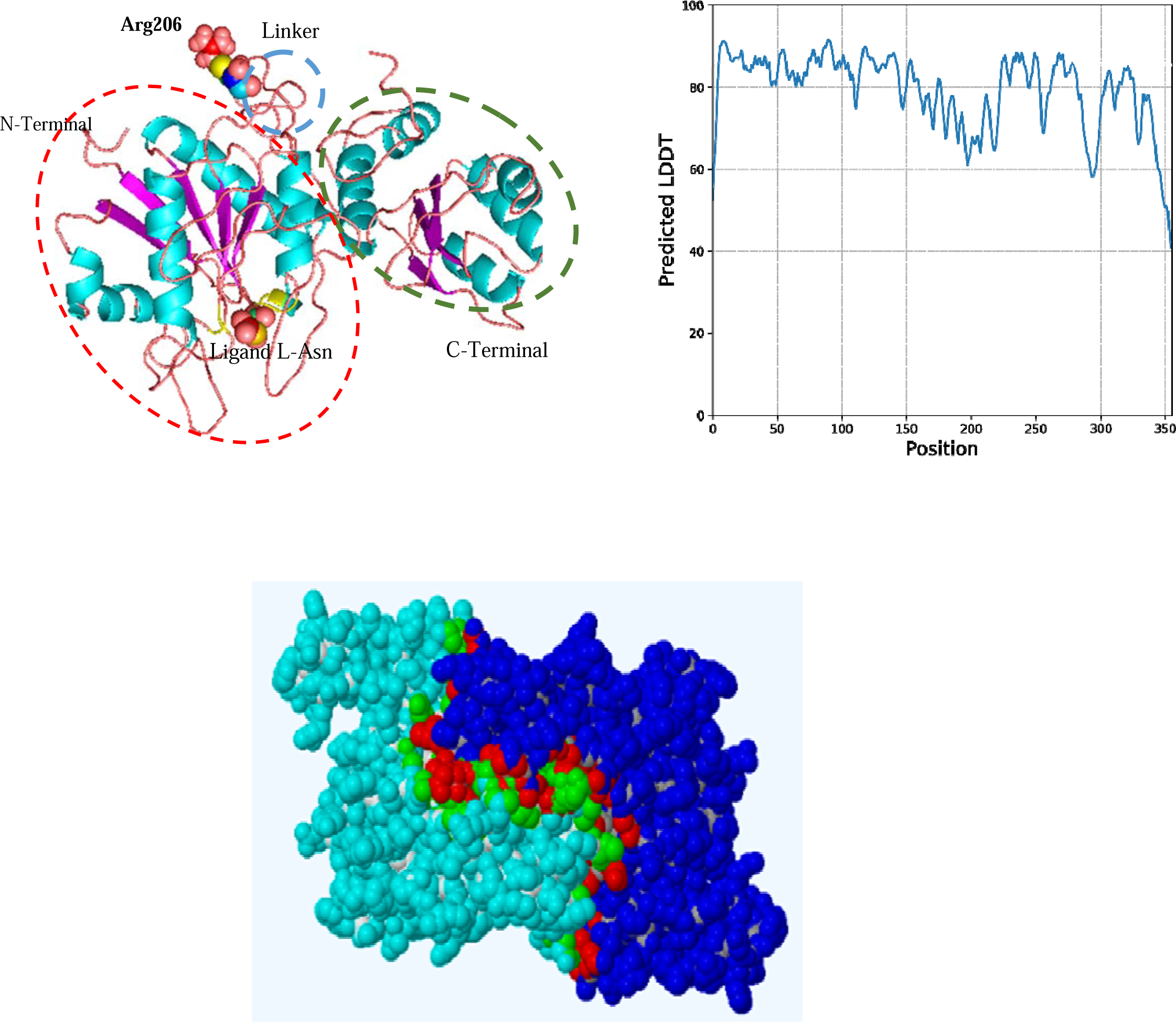
(A): The predicted 3D structure of *H. elongata* L-ASNase monomer. The larger N-terminal domain in green and the smaller C-terminal domain in blue. (B): Predicted 3D structure of *H. elongata* L-ASNase using the trRosetta server, assessed by the Local Distance Difference Test (LDDT). (C): Amino acids involved in the interaction between the two chains are highlighted in red and green by the COTH online server.

**Fig. 3.**
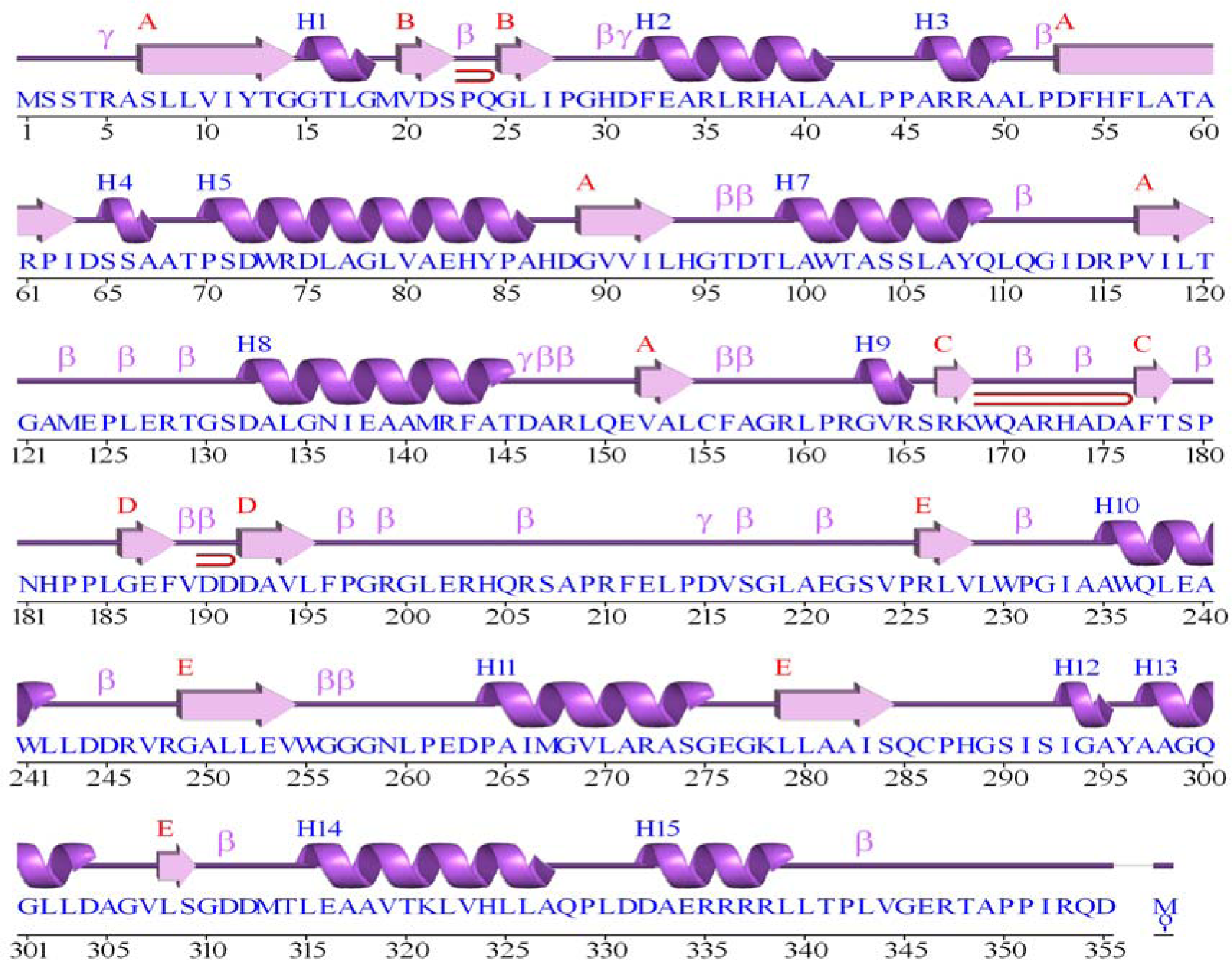
Prediction of the secondary structure of the modeled enzyme conducted using the PSSpred online server.

The Galaxy web server identified active site residues, notably Thr16, Ser65, Thr96, Asp97, and Ala122, within the modeled enzyme. These findings corroborate with results from both multiple sequence alignment and the PPSS server, highlighted in yellow in Fig. 4A. Moreover, analysis of the docking data demonstrated the formation of hydrogen bonds between the predicted active site residues and the substrate L-Asn, confirming their crucial role in substrate binding (Fig. 4B). The optimal ligand docking position concerning the substrate L-Asn and the modeled L-ASNase revealed a binding energy of −8.1 kcal/mol. The substrate binding pocket is within the N-terminal domain and involves various types of interactions. Upon observation of the correlations among residues Thr16, Ser65, and Asp97 of the modeled L-ASNase with L-Asn as the substrate, it was evident that they align in a configurational manner, resulting in a triad formation. This alignment suggests a possible catalytic function in the interaction between enzymes and substrates.

**Fig. 4.**
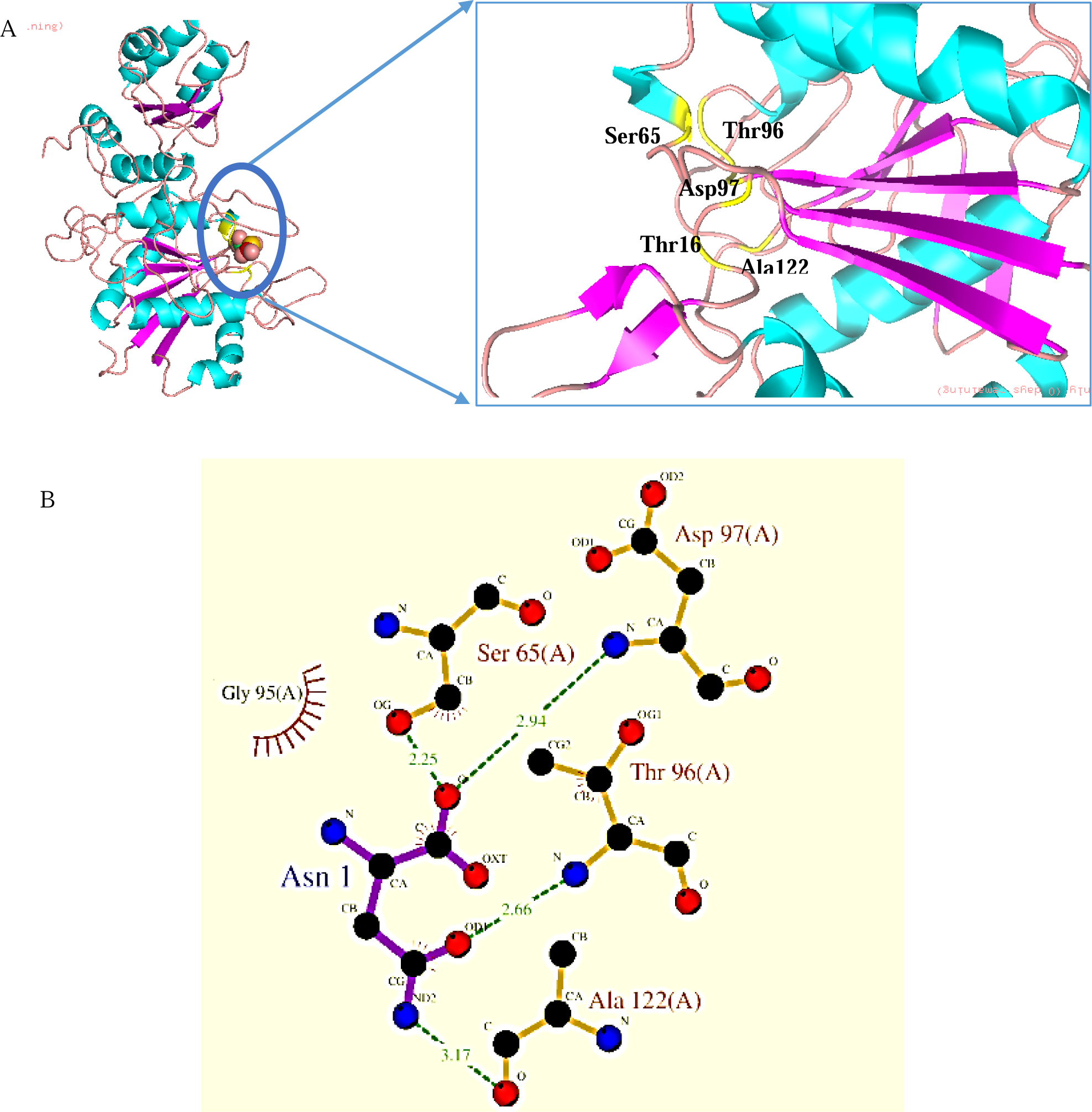
(A): Molecular docking results of L-ASNase and L-Asn conducted using the Galaxy web server. (B): LigPlot representation illustrating the interactions between the enzyme L-ASNase and the substrate L-Asn. Dashed green lines indicate the lengths of hydrogen bonds formed during the interaction.

### 3.2. Development of Mutations and Smart Libraries Guided by Sequence Input Data

To pinpoint protease-sensitive cleavage sites within the amino acid sequence of *H. elongata* L-ASNase, the PeptideCutter webserver was employed, revealing 29 predicted cleavage positions for trypsin (Fig. 5A). Notably, among the 29 identified sites, six residues, Arg172 (87%), Arg206 (69%), Arg210 (91%), Arg246 (63%), Arg353 (86%), exhibited the highest accessible surface area and are situated in the loop regions (Fig. 5B).

**Fig. 5.**
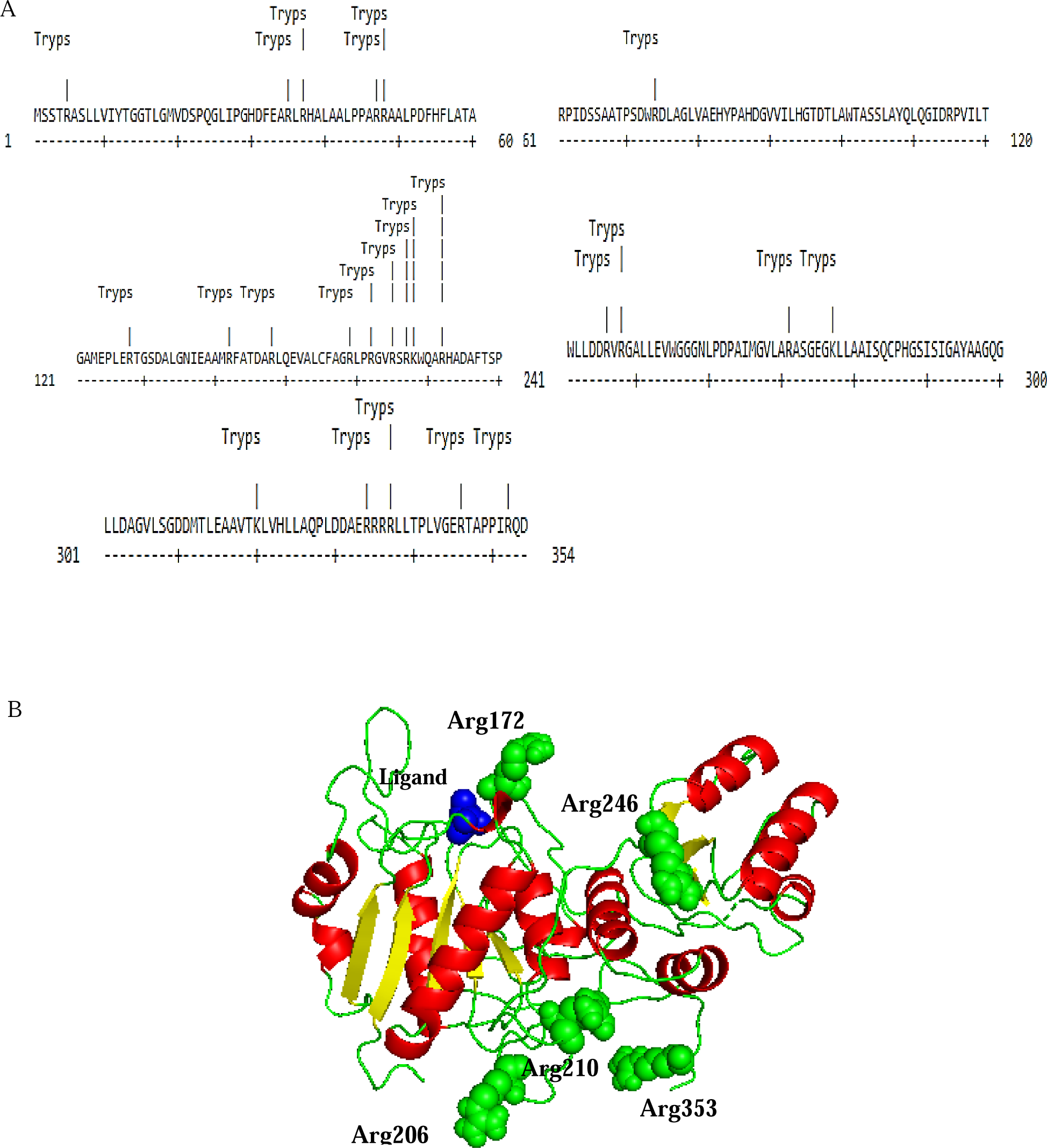
(A): Putative proteolytic map of *H. elongata* L-ASNase peptide fragments generated by the PeptideCutter following cleavage by trypsin. (B): Visualization of accessible surface areas of residues at cleavage sites of trypsin using PyMOL.

The Hotspot Wizard identified Arg206, Arg203, His204, Gln205, and Asp114 as hotspot residues for mutation in *H. elongata* L-ASNase. Among these, Arg206, located within a flexible loop region responsible for connecting the two protein domains, attained the highest mutability score of nine. Additionally, the LBtope server identified this residue as an immunogenic site in *H. elongata* L-ASNase. The computational engineering findings indicate that Arg206 may be a suitable candidate for site-saturated mutagenesis, given its positioning within the loop region, lack of proximity to the active site and non-involvement in quaternary structure formation. Notably, it is among the sites prone to trypsin cleavage, exhibiting considerable solvent accessibility. Thus, preserving the integrity of Arg206 through mutation would likely maintain the protein’s structure, stability, and activity without alteration (Fig. 5B).

### 3.3. Saturation Mutagenesis and Library screening

Two primers were employed in the PCR reaction: a forward mutagenic primer containing NNK degeneracy at the mutation site (32 codons that encode all 20 amino acids and one stop codon), and a reverse non-mutagenic primer with pET21a-*asp* as the template. Agarose gel analysis revealed a 6500 bp PCR product (Fig. S3). Following ligation, the PCR products were introduced into *E. coli* XJb (DE3) cells that are capable of autolysis. To achieve saturation mutagenesis of NNK degeneracy, 96 colonies were necessary to ensure 95% coverage of residue randomization. Five colonies exhibiting equivalent or superior specific enzyme activity compared to the wild-type were chosen for sequencing. Among them, variants Arg206Gln, Arg206Cyc, and Arg206Thr were identified, with Arg206Thr exhibiting the highest enzyme activity and stability at 37 °C, thus selected for further examination.

The specific activity of the purified wild-type and mutant enzymes was measured 1748 and 2400 U/mg, respectively. The impact of mutation on the enzyme’s affinity for the substrate and catalytic activity was investigated by calculating kinetic parameters. Utilizing the Michaelis-Menten equation with L-Asn as the substrate (Fig. S4) (Table 1).

**Table 1.** Kinetic parameters of the wild-type and mutant L-ASNases with L-Asn as substrate.

| Enzymes | Specific activity (U/mg) | K<sub>m</sub> (mM) | V<sub>max</sub> (μmol/min) |
| --- | --- | --- | --- |
| Wild-type | 1748±105 | 0.535±0.041 | 0.23±0.015 |
| Arg206Thr mutant | 2400±144 | 0.629±0.038 | 0.253±0.018 |

### 3.4. Effect of Temperature, pH, and NaCl Concentration on Enzyme Activity

Both the mutant and wild-type enzymes exhibited the maximum L-ASNase activity at 37°C (Fig. S5 A). The optimum pH for both enzymes was defined to be at pH 7.50, although the wild-type enzyme exhibited slightly higher activity at pH 8 (Fig. S5 B). The findings demonstrated that both enzymes maintained their activity at a 0.15 M NaCl, equivalent to human blood serum osmolality (Fig.S5 C).

### 3.5. Enzymes Stability Assessment Toward Trypsin and Blood Serum

The structural stability of both the mutant and wild-type enzymes under trypsin-induced proteolytic cleavage revealed that the mutant L-ASNase displays significantly improved resistance relative to the wild-type enzyme. In the case of the wild-type enzyme, activity decreased by approximately 46% and 55% after 90-120min, respectively, with a trypsin to enzyme ratio of 0.5%. Notably, the wild-type enzyme was largely denatured after 120min under these conditions (Fig. 6A). Conversely, analysis of the mutant enzyme’s resistance against trypsin shown that even after 90-120min of incubation with a 5% trypsin to enzyme ratio, it retained 62% and 45% of its activity, respectively (Fig. 6B). This decline in activity was corroborated by SDS-PAGE results (data not shown).

**Fig. 6.**
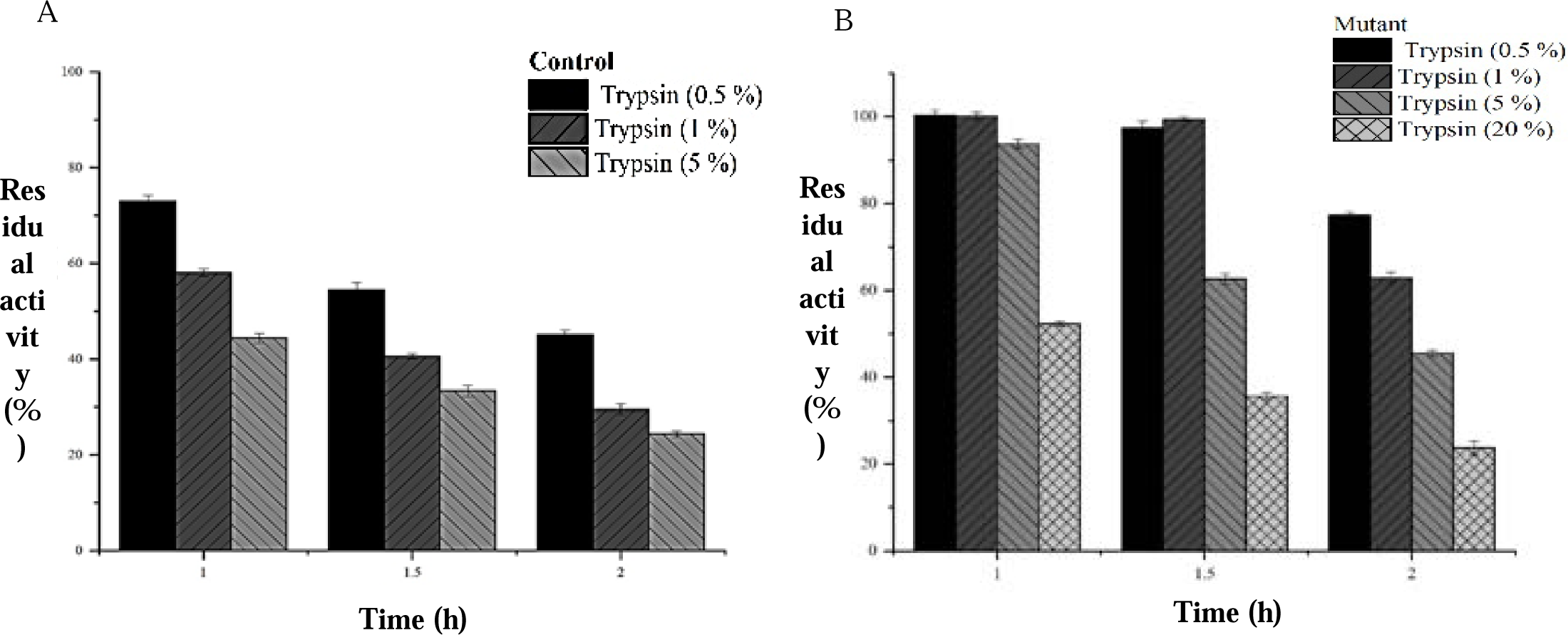
Residual activity over time during trypsin digestion of (A) the wild-type and (B) the mutant enzyme.

The half-lives of both mutant and wild-type enzymes were determined by incubating them with human serum at 37°C, followed by estimating the decay curve fitting using a one-phase model. The wild-type enzyme exhibited a half-life of 1.8h, whereas the mutant enzyme had a half-life of 3.47h (Fig. 7A). In the case of phosphate buffer instead serum, both enzymes retained their activity within the reaction mixture, with a half-life of 2.26h for the mutant and 2.33h for the wild-type (Fig. 7B).

**Fig. 7.**
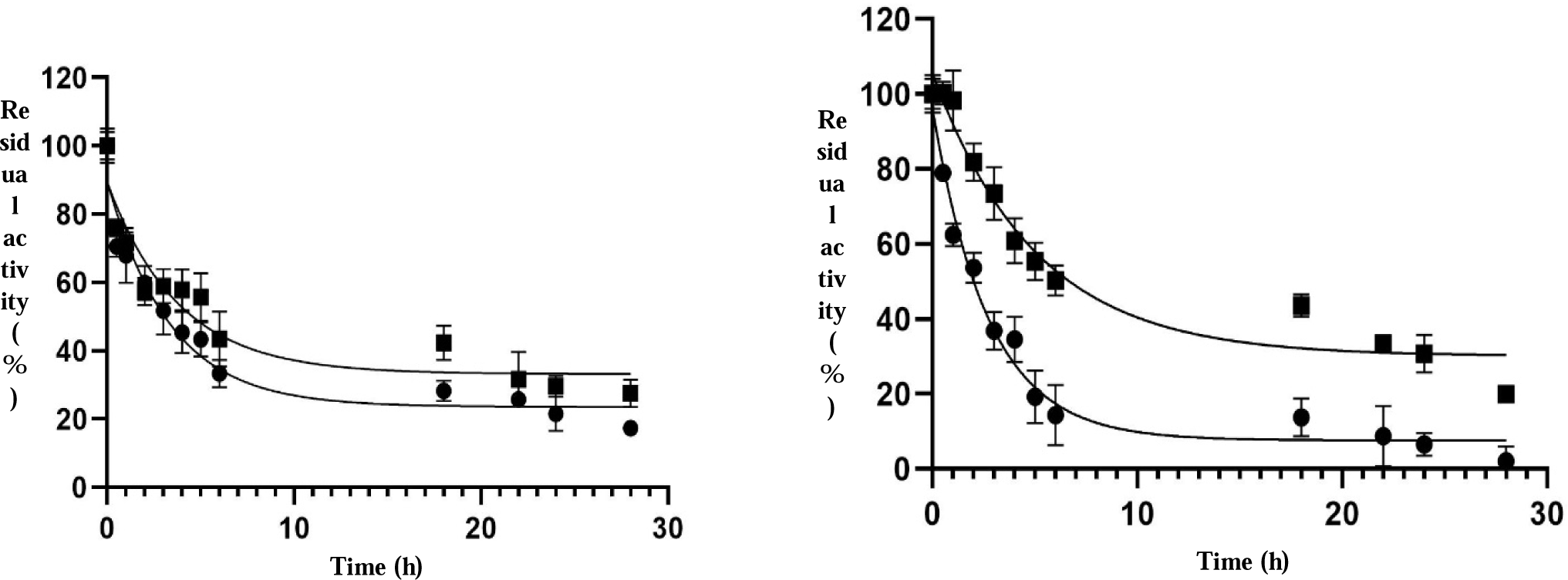
(A): Half-life comparison of the wild-type and mutant enzymes in human serum. (B): Half-life comparison of the wild-type and mutant enzymes in phosphate buffer at 37 °C. Wild type (●), Mutant (▪)

### 3.6. Enzyme Cytotoxicity Assessment

The antiproliferative efficacy of wild-type and mutant *H. elongata* L-ASNase was assessed on the K562 cell line via the MTT assay. Following a 24h incubation period, increasing concentrations of both enzymes led to a dose-dependent reduction in cell viability, significantly lowering cell survival rates compared to controls. Our findings reveal approximately equal IC_50_ values of 1.54±0.098 U/mL and 1.45±0.083 U/mL for wild-type and mutant L-ASNase, respectively. In a hemolytic test on human blood, neither enzyme displayed hemolytic activity at concentrations up to 6 U/mL, contrasting with the positive control containing 1% SDS lysis solution. Additionally, assessment of cytotoxicity on non-leukemic HUVEC cell lines via the MTT assay demonstrated no adverse effects at concentrations up to 6 U/mL. These findings confirm the robust cytocompatibility of both wild-type and mutant enzymes (Fig. 8).

**Fig. 8.**
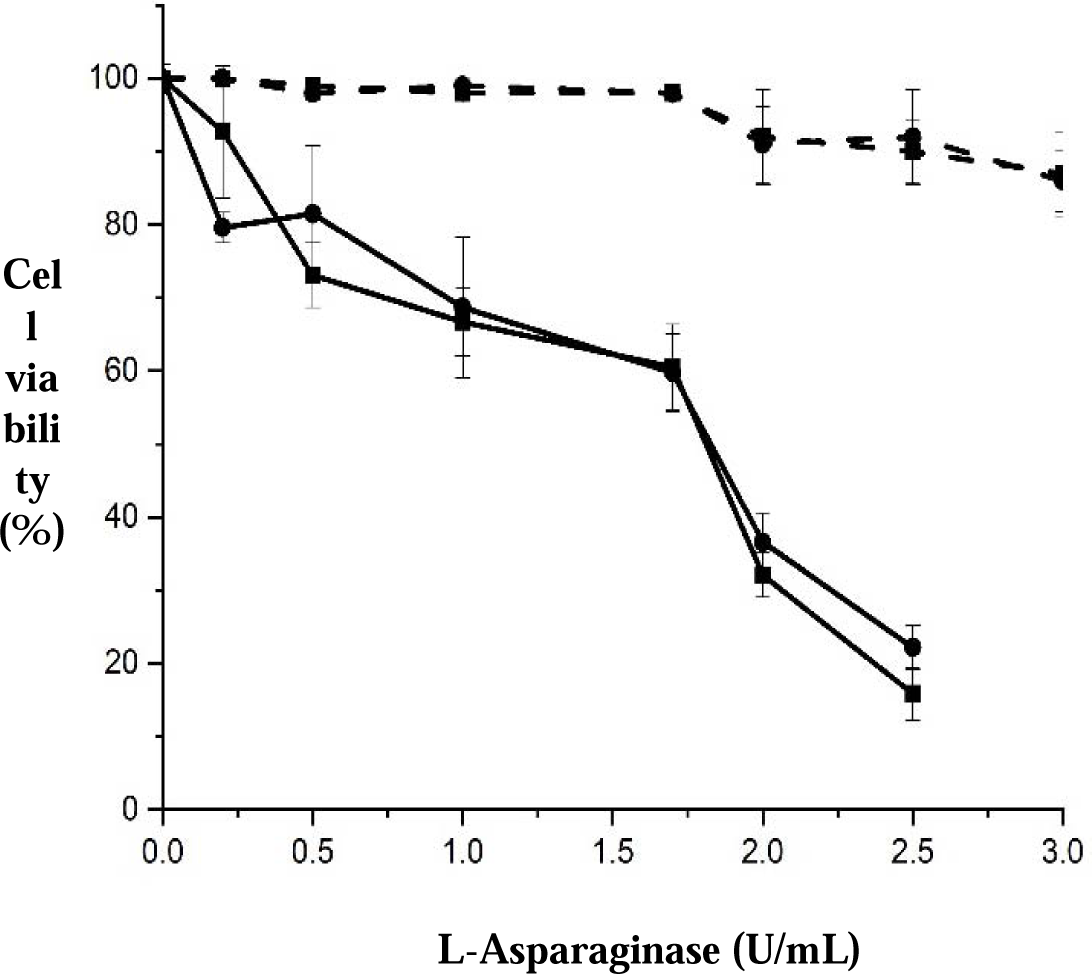
(A): Assessment of the anti-proliferative effects of wild-type and mutant *H. elongata* L-ASNase on the K562 and HUVEC cell lines. Wild type (▪) and Mutant (●), HUVEC cell line (dashed line) and K562 cell line (solid line)

According to DynaMut analysis, the substitution of Arg with Thr at position 206 resulted in a ΔG value of −0.254 kcal/mol, indicating a favorable effect on protein stability (Fig. 9). Additionally, the calculation of vibrational entropy energy for the mutant yielded −0.133 kcal/mol, indicating a reduction in molecular flexibility.

**Fig. 9.**
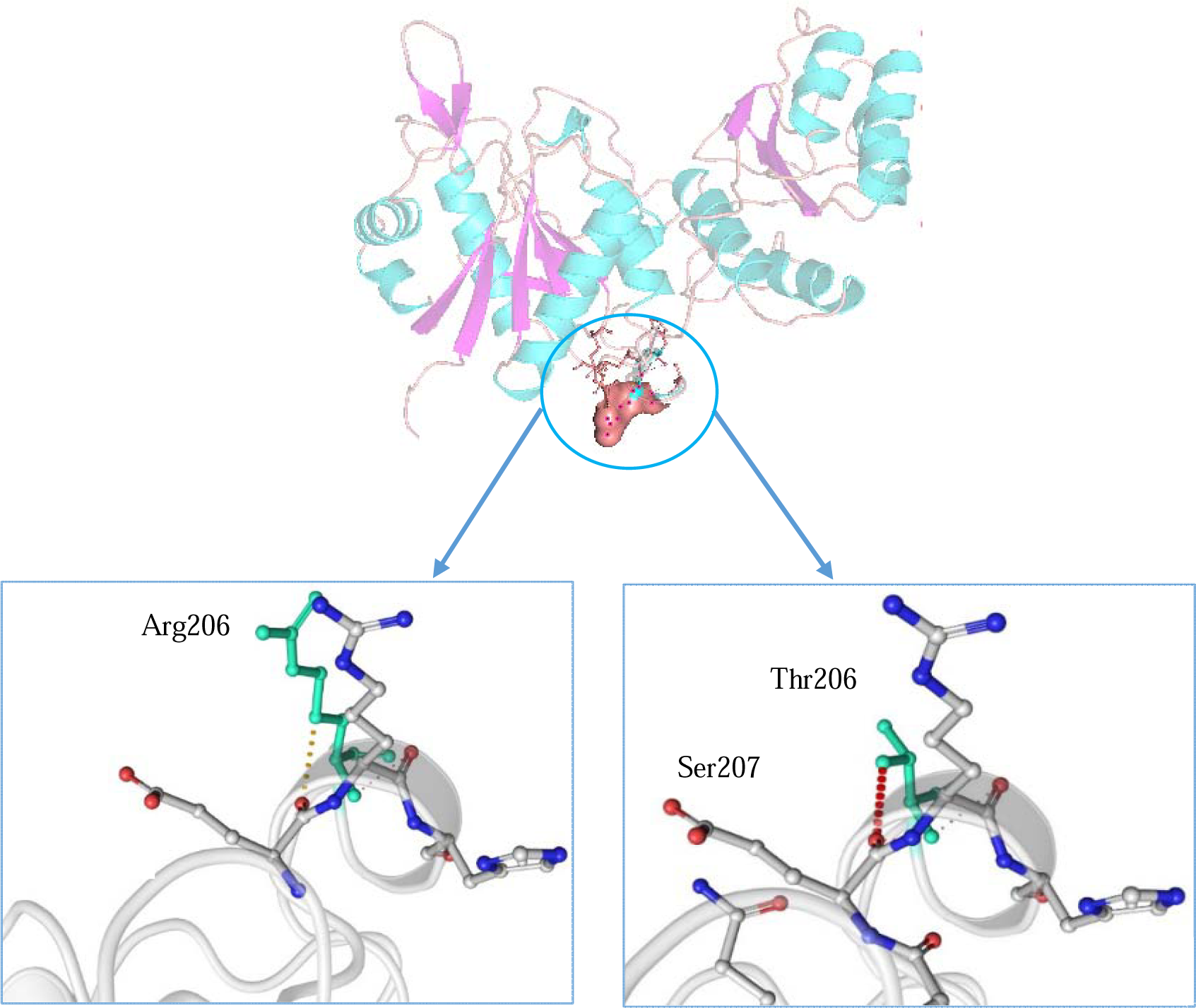
Residues involved in interactions, including wild-type and mutant residues along with their surrounding residues, are depicted in light-green color. (A): Wild-type. (B): Mutant.

The Residue Interaction Network Generator (RING) online server analyzed the *H. elongata* L-ASNase structure, identifying 254 hydrophobic interactions, 5 ionic bonds, and 268 hydrogen bonds. In the protein-ligand complex, 5 hydrogen bonds were predicted. Comparative analysis using RING showed that the mutant enzyme exhibited increased hydrogen bonds (271) and van der Waals bonds (255) compared to the wild-type, suggesting enhanced stability. Additionally, surface accessibility analysis indicated reduced exposure of the Thr residue at position 206 and its surrounding residues relative to the wild-type residue (Fig. S6 A, B). Arg206Thr substitution has also modified the epitope map, leading to a smaller epitope fragment compared to the wild-type. Furthermore, the surface area occupied by Thr in the epitope map has decreased in comparison to the wild-type.

## 4. Discussion

Asparaginase therapy is crucial in acute lymphoblastic leukemia treatment, but its clinical use is limited by challenges like immune responses, instability in the bloodstream [35–37]. Commercial drugs may trigger allergic reactions and lose effectiveness due to protease degradation [38]. Protein engineering offers promising solutions to enhance the stability of therapeutic proteins while minimizing side effects and treatment costs. [39–40]. Leveraging halophilic bacteria, we aim to enhance the proteolytic stability of *H.elongata* L-ASNase while maintaining its catalytic activity and structural integrity, utilizing innovative computer modeling to identify mutation sites for improved enzyme stability.

Through sequence alignment, it shares %. 37% identity with *Y. pestis* L-ASNase (PDB accession no. 3NTX), 37.1% identity with *E. coli* type I L-ASNase (PDB accession no. 3C17), and 22.1% with *E. coli* type II (PDB accession no. 3ECA). This comparative analysis highlights both conserved and divergent regions across species, shedding light on potential functional differences and evolutionary adaptations. Utilizing the trRosetta server, we modeled its tertiary structure, achieving a high TM-score of 0.98, indicating structural accuracy. Subsequent analyses, including root-mean-square deviation (RMSD), Ramachandran plot analysis, and ERRAT evaluation, affirmed the geometric fitness, stereochemical quality, and resolution of the modeled structure [41]. Exploring the functional attributes of *H. elongata* L-ASNase, we delineated key residues constituting its binding site. This analysis revealed a cluster of amino acids, including Thr16, Ser65, Thr96, Asp94, and Ala122, distributed within the N-terminal domain’s flexible loop region. The residues displayed a significant level of conservation among various species, suggesting their crucial function in catalytic activity and substrate recognition. Moreover, comparing the predicted binding site with previously characterized L-ASNase variants underscored similarities in substrate interaction patterns, particularly with *E. coli* and *Bacillus subtilis* counterparts, albeit with varying binding free energies (41). The active site relies on a conserved Thr residue acting as the primary nucleophile, with a predicted strong affinity for L-Asn (−8.1 kcal/mol) [42]. The secondary structure analysis of *H. elongata* L-ASNase revealed a composition of 42% random coils, 33% alpha helices, and 20% extended strands, mirroring the insights gained from circular dichroism (CD) data [27]. *H. elongata* L-ASNases modeled structure comprises two domains, with a dimeric architecture similar to *Y. pestis*, suggesting interaction between chains involving approximately 50 residues. The proteolytic mapping of wild-type L-ASNase revealed 29 trypsin recognition sites, with six residues exhibiting the highest solvent accessible surface located in flexible regions. Using the HotSpot Wizard and LBtope analysis, we identified Arg206 as an immunogenic hotspot site, located in a flexible loop distant from the active site, prompting its selection as the mutation site. Through site-saturated mutagenesis, the Arg206Thr mutant of *H. elongata* L-ASNase was created, showing enhanced stability against trypsin, human serum and increased specific activity with insignificant change in *Km* value, in comparison to the wild-type enzyme. Both mutant and wild-type enzymes exhibited significantly higher specific activities and serum stabilities compared *to E. chrysanthemi* and *E. coli* L-ASNases [43]. Furthermore, optimal activity for both variants was at 37 °C and pH 7.50, mirroring physiological conditions. No adverse effects were observed on the HUVEC cell line and red blood cells when either of the enzymes was present, indicating their non-cytotoxic nature. However, these enzymes effectively suppressed the growth of cancer cells, demonstrating their selective inhibitory effects. The mutant Arg206Thr exhibited altered interatomic interactions, leading to increased stability and reduced flexibility of its structure due to the formation of a network of weak hydrogen bonds, augmented hydrophobic interactions, decreased protein surface charge, and reduced accessibility. The substitution of Thr, a polar but neutral amino acid, for Arg, which carries a positive charge, results in the replacement of ionic bonds with hydrogen bonds. Moreover, weak hydrogen bonds are formed between Thr206 and surrounding residues (Ser207 and His204) in the mutant state. The mutant Arg206Thr also induces changes in the epitope map and reduces surface accessibility. Previous studies have established a direct link between protein thermodynamic stability and proteolytic sensitivity, with protein dynamics playing a crucial role in structure stability and proteolytic susceptibility [44–50].

## 5. Conclusion

In sum, the application of a semi-rational technique in the engineering of *H. elongata* L-ASNase has proven to be successful in achieving proteolytic resistance and enhancing enzymatic activity. This represents the first instance of designing proteolytic resistance in *H. elongata* L-ASNase. These findings underscore the potential of *H. elongata* L-ASNase as a therapeutic agent, offering improved stability and compatibility with serum physiochemical characteristics, thereby reducing the need for frequent injections.

### Declaration of Generative AI and AI-assisted technologies in the writing process

In the course of preparing this work, the authors made use of the https://www.editmyenglish.com/freeediting.aspx service and ChatGPT to verify the grammatical correctness, spelling accuracy, and fluency of the English writing.

## Acknowledgement

The authors would like to express their gratitude to the Iran National Science Foundation (INSF) for providing financial assistance for this research project (4004751).

